# Comparative Transcriptomics Reveals Inflammatory and Epigenetic Programs that Actively Orchestrate Pineal Brain Sand Calcification

**DOI:** 10.1101/2025.06.16.660032

**Authors:** Giovanni J.A. Vazquez Ramos

## Abstract

**Background:** The pineal gland secretes melatonin but paradoxically calcifies more than any other intracranial structure, forming hydroxyapatite “brain-sand” (corpora arenacea) that correlates with reduced melatonin output, sleep disruption and heightened neuro-degenerative risk. Whether this mineralization is a passive dystrophic event or an active, bone-like process remains unclear.

**Methods:** Analyzed RNA-seq datasets from pineal glands of six vertebrate species, calcifiers *Homo sapiens*, *Rattus norvegicus,* and *Capra hircus*, versus non-calcifiers *Mus musculus*, *Gallus gallus* and *Danio rerio*. Species-specific transcripts were mapped to human orthologues, merged, and filtered. Phylogenetically informed differential-expression testing used Brownian-motion and Pagel’s λ phylogenetic generalized least-squares models, calibrated on a TimeTree divergence phylogeny. Genes significant in both models (|log₂FC| > 1; FDR < 0.05; λ < 0.7) were assigned to functional pathways and visualized by PCA, heat-mapping and volcano plots.

**Results:** Calcifying species segregated cleanly from non-calcifiers on the first two principal components, reflecting a shared 103-gene “calcifier module”. Top up-regulated transcripts included developmental morphogens (*GLI4, IQCE, NOTCH4*), epigenetic regulators (*SETD1A, ZNF274, ATF7IP*), inflammatory mediators (*CSF2RB*), and quality-control factors (*GABARAPL2, RHOT2*). Every leading candidate exhibited minimal phylogenetic signal (λ ≈ 0), indicating that differential expression tracks the calcified phenotype rather than shared ancestry. Conversely, only three genes (*RMI2, RASL11B, GPR18*) formed a non-calcifier module, suggesting potential protective roles that are down-regulated during mineralization.

**Conclusions:** Pineal calcification is not a passive by-product of aging but a regulated, lineage-restricted program that redeploys Hedgehog, Notch and chromatin-remodeling pathways classically required for skeletal ossification. The ten-gene core signature identified offers a molecular foothold for mechanistic dissection and therapeutic targeting aimed at preserving pineal function and circadian health.

**Significance:** This is the first phylogenetically controlled transcriptomic survey to link pineal “brain-sand” formation to specific developmental and inflammatory gene networks, revealing convergent evolution of calcification programs across divergent mammalian lineages.

## Introduction

The pineal gland is a unique neuro-endocrine structure that translates photic information into the rhythmic secretion of melatonin, a pleiotropic hormone with antioxidant, circadian-entraining and neuroprotective properties.^1^ Despite its small size, the gland calcifies more frequently than any other intracranial tissue: layered concretions of hydroxy-apatite-like calcium/magnesium salts (corpora arenacea, “brain-sand”) are detectable on computed tomography in more than half of the adult population, with a pooled prevalence of ≈ 62 % in contemporary meta-analysis.^2^ Histological surveys dating back to the 20th century confirm that the number and diameter of acervuli increase markedly after puberty and into senescence, yet remain scant or absent in many non-mammalian vertebrates.^3^ While often dismissed as an epiphenomenon of aging, pineal calcification correlates with reduced melatonin output, sleep-wake disruption and heightened risk of neuro-degenerative disorders including Alzheimer’s disease, underscoring the need to decipher its molecular origins.^4^ Notably, wild-type mice exhibit negligible pineal calcification, making them ideal negative-control models for comparative studies of mineralization. ^5^ In contrast, rats develop extensive pineal concretions, approaching human-like rates by late adulthood, underscoring their utility as positive-control models for investigating the mechanisms and consequences of glandular calcification.^6^

### Passive precipitation or programmed ossification

Early ultrastructural work demonstrated that hydroxy-apatite first accumulates inside degenerating dark pinealocytes, with mineral foci “sometimes associated with cellular debris,” a pattern typical of dystrophic calcification.^7^ Yet several observations argue against a wholly passive process. First, human concretions exhibit multi-layered concentric lamellae and plate-like crystals “similar to dentin and bone”.^8^ Second, comparative surveys show striking species variation: rats and humans calcify readily, whereas mice and several other mammals form few or no acervuli throughout life.^3^ Third, prenatal hypoxia, a recognized inflammatory stressor, significantly increases the burden of calcium-rich particles in rat pineal glands, indicating that calcification can be experimentally accelerated independent of age.^9^ Recent molecular studies add further support: single-gene analyses have identified proteins such as retinoschisin, which are “required for pineal gland calcification and the architecture of mineral deposits” in rodents.^10^ Yet these studies could not disentangle gene–calcification relationships from phylogenetic relatedness, leaving open whether observed signatures were cause, consequence or mere species artifacts, a gap that directs attention to the signaling pathways that may actively seed calcification.

### Developmental and inflammatory cues as candidate drivers

Biological parallels between ectopic bone formation in vascular disease and corpora arenacea have fueled the view that pineal calcification is an active, bone-like process rather than a passive precipitation event.^1^ Hedgehog signaling, acting through GLI transcription factors, is essential for driving mesenchymal progenitors toward Runx2⁺ osteoblasts during normal skeletogenesis.^11^ Pathological over-activation of the same pathway has been identified as a key driver of heterotopic ossification in soft tissues.^12^ In the vasculature, Notch1 signaling, alone or in concert with the osteogenic morphogen BMP-2, can induce Msx2 expression, alkaline-phosphatase activity and matrix mineralization in vascular smooth-muscle cells, processes central to atherosclerotic calcification.^13^ Whether comparable Hedgehog or Notch-centered programs are redeployed in the aging pineal gland, and, if so, which downstream effectors or stress pathways nucleate the first mineral seeds, remains to be elucidated.

### Knowledge gaps and study rationale

To date, molecular work on pineal calcification has been fragmentary: reviews still note that “the detailed mechanisms … are not fully understood,” and the few bench studies that exist typically examine one species and a handful of genes, for example, the recent discovery that Retinoschisin alone can govern calcification in mice.^4,10^ A systematic, evolution-conscious approach is therefore missing. Importantly, several vertebrate lineages (e.g. mouse, chicken, zebrafish) retain a relatively non-calcifying pineal gland despite sharing fundamental melatonin circuitry, providing a natural experiment to distinguish causal from incidental expression changes.^3,5^ Leveraging this comparative framework could pinpoint core gene networks that are necessary, and perhaps sufficient, for the calcifying phenotype.

## Methods

### RNA-Seq Data Acquisition and Preparation

RNA sequencing datasets from pineal glands across six vertebrate species, Human (*Homo sapiens*), Rat (*Rattus norvegicus*), Mouse (*Mus musculus*), Zebrafish (*Danio rerio*), Chicken (*Gallus gallus*), and Goat (*Capra hircus*), were retrieved from the NCBI Sequence Read Archive (SRA) (accessions: SRR5756462, SRR827499, SRR290898, SRR592702, SRR6116774, SRR24843922).^14–19^ Pineal gland calcification status for each species was assigned based on existing literature.^1,3^

Transcript-level quantifications were generated using the Salmon software, and counts were summarized at the gene-level using the tximport package.^20,21^ Transcript-to-gene mappings were constructed via biomaRt with the Ensembl datasets for each species.^22^

### Cross-Species Ortholog Mapping

Orthologous genes across species were mapped to human gene symbols to facilitate direct comparisons. Initially, species-specific Ensembl IDs were mapped to human orthologs using the `orthogene` package.^23^ For species-specific datasets lacking direct orthology via g:Profiler, a fallback approach using `biomaRt` was implemented to ensure robust ortholog mapping.^22^

### Data Integration and Filtering

Gene expression matrices for each species were merged based on shared human gene symbols. Genes with zero counts across all species were removed, resulting in a filtered dataset containing only genes expressed in at least one species.

### Phylogenetic Analysis

To control for phylogenetic relationships among species, a phylogenetic tree was constructed based on divergence times obtained from TimeTree5.^24^ Phylogenetic generalized least squares (PGLS) regression was conducted using the phylolm package to test the relationship between gene expression and pineal calcification status.^25^ Both Brownian motion (BM) and Pagel’s lambda models were applied.

### Statistical Analysis and Visualization

Genes significantly associated with calcification (adjusted p-value < 0.05; log2 fold-change > 1) under both BM and lambda models (lambda < 0.7) were identified. Adjustments for multiple comparisons employed the Benjamini-Hochberg correction.

Heatmaps, PCA, volcano plots, and phylogenetic trees were generated using pheatmap, ggplot2, gridExtra, ape and ggtree.^26–30^ Visualization facilitated interpretation of gene expression patterns associated with pineal calcification.

### Data Availability

RNA sequencing data are publicly available in the Sequence Read Archive (SRA).

Source code and documentation is available under GPL v3.0 license at https://github.com/gvazram/BrainSand

## Results

### Phylogenetic distribution of pineal calcification

Mapping pineal acervuli onto a time-calibrated chronogram of six vertebrates (one specimen per species) highlights a strikingly patchy distribution: mineral deposits are observed only in Rattus norvegicus, Capra hircus and Homo sapiens, whereas Mus musculus, Gallus gallus and Danio rerio remain entirely uncalcified (Fig. 1). The tree, oriented from the present at the tips (0 Ma, right) toward the ancestral root (∼430 Ma, left), is calibrated using divergence estimates from TimeTree 5.0. Major nodes include the teleost–amniote split at ≈429 Ma, the bird–mammal divergence at ≈319 Ma, the ruminant–primate/rodent split at ≈94 Ma and the recent mouse–rat separation at ≈11.6 Ma.

**Figure 1.**
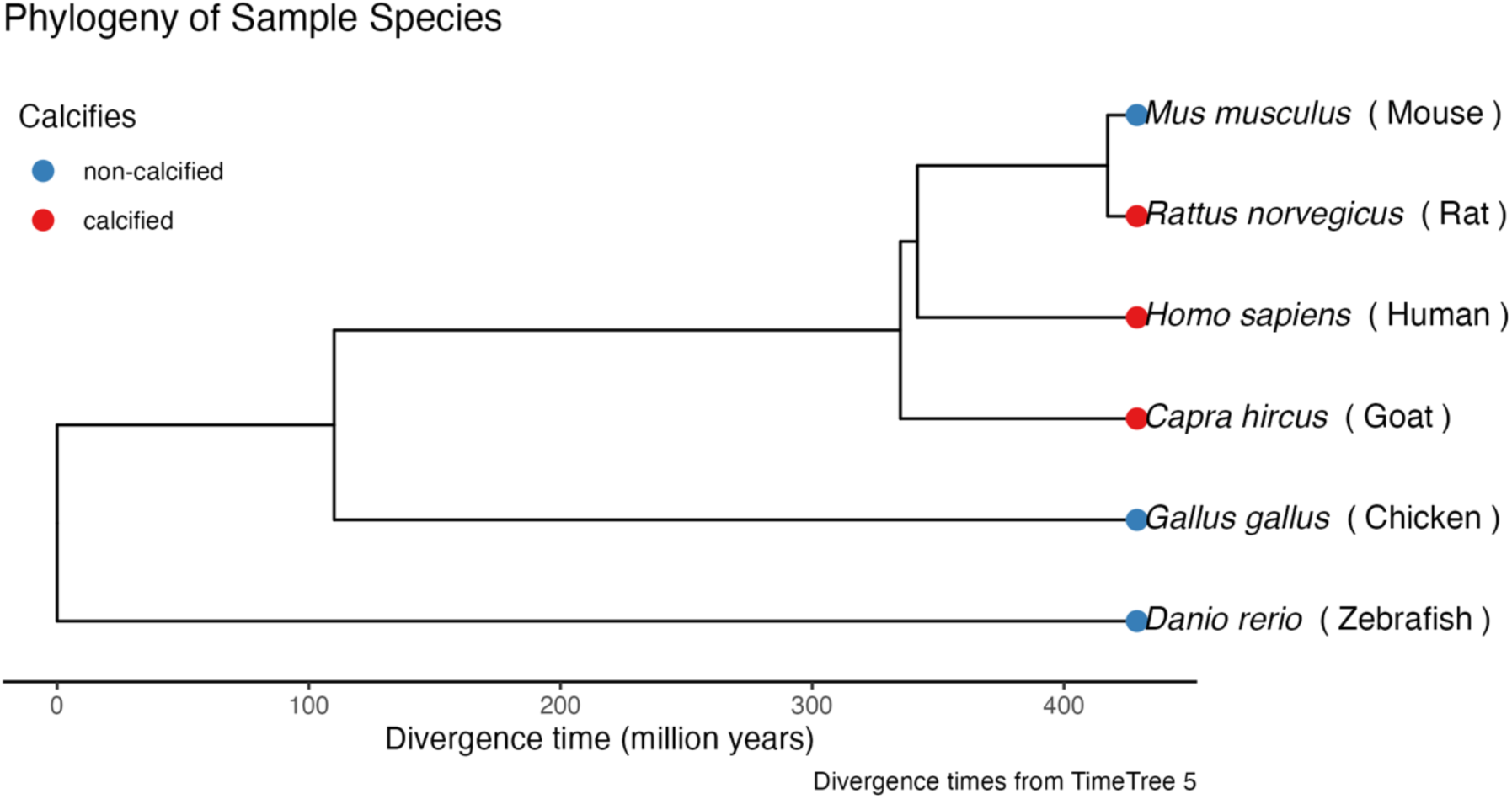
A time-calibrated phylogeny of six vertebrate species reconstructed using divergence times from TimeTree 5 shows that *Danio rerio* (Zebrafish; blue point) diverged from the amniote lineage at ≈ 429 Ma, *Gallus gallus* (Chicken; blue) split from the mammalian line at ≈ 319 Ma, and within mammals *Capra hircus* (Goat; red) branched off at ≈ 94 Ma; the primate lineage (*Homo sapiens*; red) and rodent lineage (*Mus musculus* and *Rattus norvegicus*) then separated at ≈ 87 Ma, with Mouse (blue) and Rat (red) themselves diverging most recently at ≈ 11.6 Ma. Tip labels combine italicized scientific names with common names in parentheses, and colored points denote calcification status: blue = non-calcified, red = calcified.

This distribution can be interpreted under two equally parsimonious scenarios. The first posits an early origin of pineal calcification on the stem leading to crown mammals, soon after the bird– mammal split, with a subsequent, lineage-specific loss in the mouse lineage. Under this model, Mus musculus would have lost the capacity to form acervuli while its sister taxon, R. norvegicus, retained it. Alternatively, pineal acervuli may have arisen independently at least twice: once on the early mammalian stem and again within Rodentia, giving rise to deposits in rat but not in mouse.

Distinct absence of calcification in both the avian Gallus and the teleost Danio further underscores that mineralization of the pineal gland is not an ancestral vertebrate trait but rather a derived feature confined to specific mammalian clades. The presence of acervuli in goat and human, separated by ∼94 Ma of independent evolution, points to convergent molecular mechanisms co-opted in each lineage. Conversely, the rodent comparison, mouse versus rat, suggests that relatively minor genetic or regulatory changes can abolish or permit pineal mineral deposition over a short evolutionary timeframe (∼11.6 Ma). Together, these phylogenetic patterns imply that pineal calcification has evolved via lineage-restricted gains and/or losses, reflecting dynamic shifts in the regulation of mineralization pathways across vertebrate history.

### Transcriptomic separation by calcification status

A principal-component analysis of log₂(TPM + 1)–transformed expression profiles across all six pineal glands reveals a striking segregation of calcifying versus non-calcifying species along the two major axes of variance (Fig. 2). On the biplot, where each point is labeled with its *italicized* scientific name and parenthesized common name, the non-calcified taxa (*Danio rerio* (Zebrafish), *Mus musculus* (Mouse), *Gallus gallus* (Chicken)) form a tight cluster in the upper-right region, reflecting a shared transcriptional program centered on canonical circadian and neuroendocrine genes. By contrast, calcifying species (*Rattus norvegicus* (Rat), *Capra hircus* (Goat), *Homo sapiens* (Human)) spread into the lower-left and far-left quadrants, indicating both a divergent core expression signature and increased heterogeneity in pathways linked to mineral deposition. Notably, *D. rerio* stands apart from its non-calcifier peers by its extreme elevation on the second component, suggesting additional zebrafish-specific regulatory shifts, while the remaining non-calcifiers occupy more intermediate positions along that same axis. This clear partitioning, despite single-replicate sampling, underscores an intrinsic transcriptional divide that aligns closely with the presence or absence of pineal concretions and highlights candidate gene sets for further functional validation.

**Figure 2.**
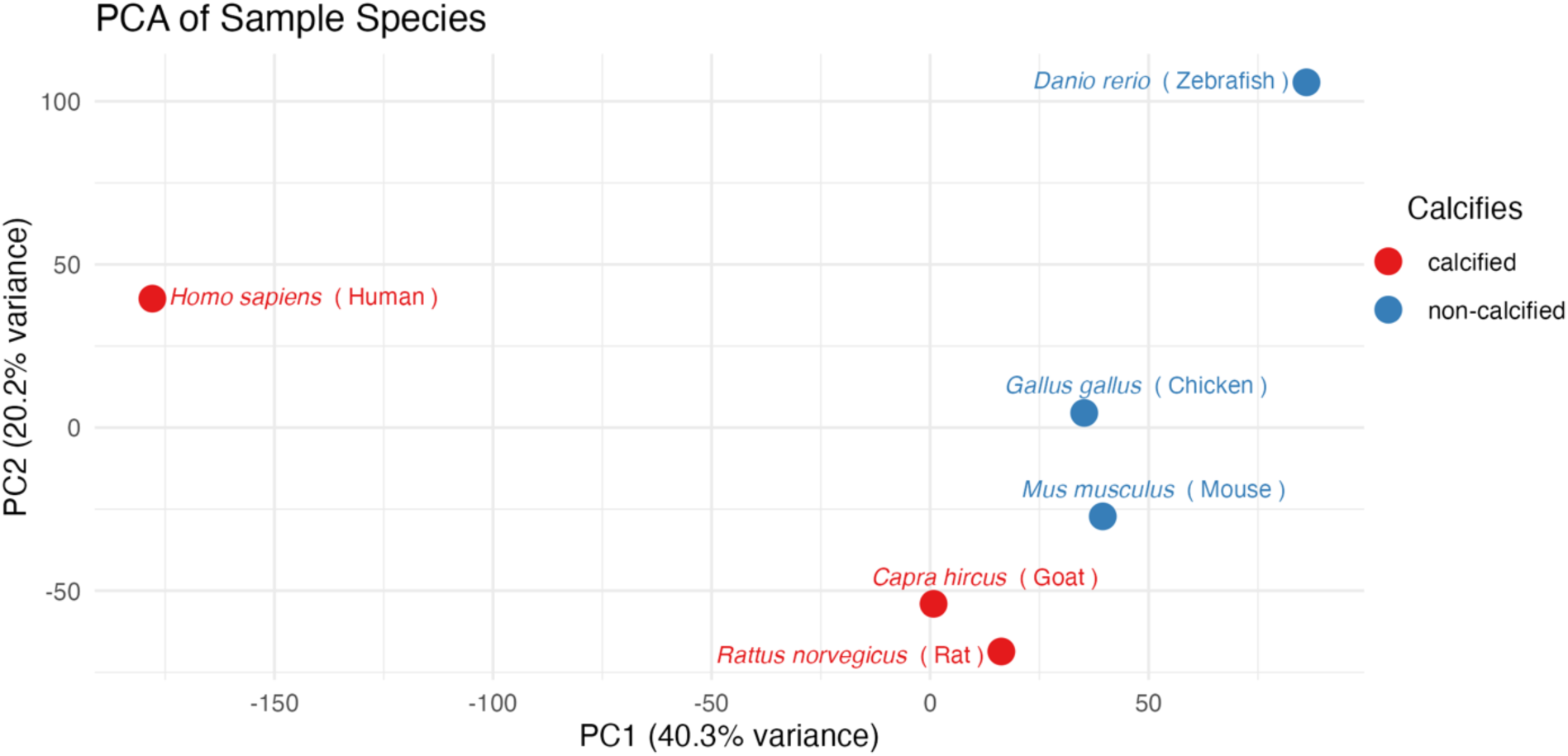
Principal-component analysis (PCA) of log₂(TPM + 1)–transformed expression profiles across six vertebrate species. Shown are the first two principal components, PC1 (40.3% of total variance) on the x-axis and PC2 (20.2%) on the y-axis, with each point labeled by its italicized scientific name and common name. Non-calcified species (blue) cluster in the upper-right quadrant, whereas calcified species (red) occupy the lower-left or far-left quadrants.

### Calcification-associated gene modules

In a PGLS-filtered set of 106 genes (adj. Pλ < 0.05) shown in Figure 3, row-scaling each transcript to a mean of zero and SD of one and mapping Z-scores to a 100-step blue–white–red palette (blue ≤ –2 = low, white = 0, red ≥ +2 = high) reveals an almost unanimous up-regulation in calcifiers (human, rat, goat) contrasted against non-calcifiers (mouse, chicken, zebrafish). Hierarchical clustering (Euclidean distance, complete linkage) partitions these genes into two discrete blocks: a vast calcifier module of 103 genes, enriched for developmental morphogens (GLI4, IQCE, NOTCH4)^31–33^, cytokine and stress-response mediators (CSF2RB, APEH)^34,35^, epigenetic writers and erasers (SETD1A, ZNF274, ATF7IP)^36–38^, autophagy/mitochondrial quality-control factors (GABARAPL2, RHOT2)^39,40^ and numerous metabolic enzymes, marked by intense red peaks in calcifying species and deep blue troughs in non-calcifiers; and a tiny non-calcifier module comprising only RMI2, RASL11B and GPR18, each of which flips the pattern, showing higher Z-scores in non-calcifiers (red) and suppression in calcifiers (blue). The presence of just these three lineage-specific outliers, RMI2, a recombination factor; RASL11B, a small GTPase regulator; and GPR18, an orphan GPCR, suggests they may play protective or housekeeping roles that are down-tuned during the active, bone-like mineralization program.^41–43^ Taken together, the sharp bimodal color gradients and matching dendrograms confirm that pineal calcification is driven by a tightly coordinated transcriptional network uniquely activated in calcifying mammals.

**Figure 3.**
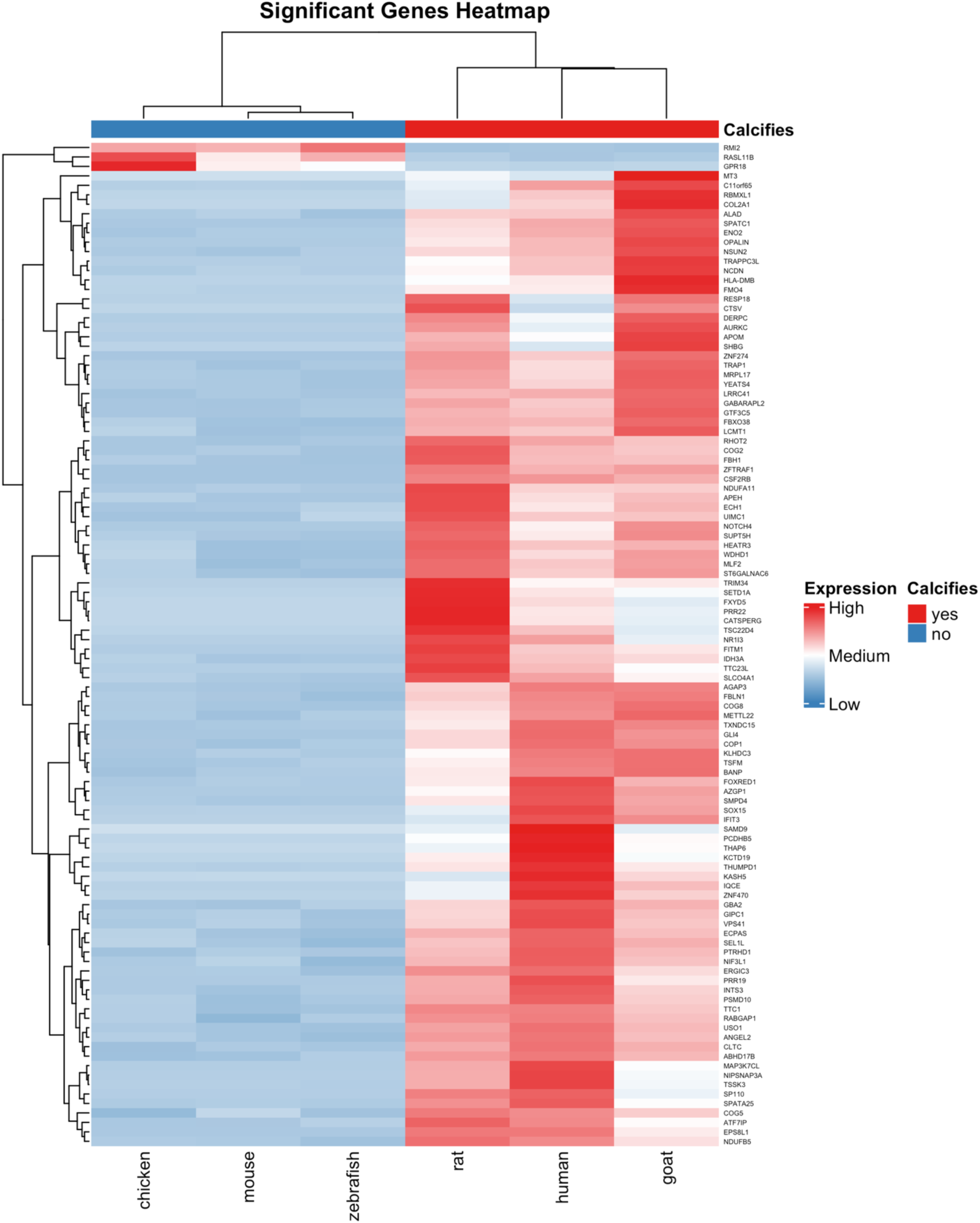
Heatmap of row-scaled (Z-score) expression for genes significantly associated with pineal calcification (PGLS adj. Pλ < 0.05) across six vertebrates: human, rat, goat (calcifiers) and mouse, chicken, zebrafish (non-calcifiers). Expression values were scaled per gene to mean 0 and SD 1, then mapped to a 100-step blue–white–red palette (blue ≤ –2 = Low, white = 0 = Medium, red ≥ +2 = High; legend ticks at –2, 0, +2). Hierarchical clustering (Euclidean distance, complete linkage) was applied to both rows (genes) and columns (species), with dendrograms shown. A discrete annotation bar above the heatmap indicates calcification status (red = yes; blue = no).

### Phylogenetically informed differential expression (volcano-plot) analysis

Phylogenetically informed volcano-plot analysis (Fig. 4) identified the top ten transcripts whose expression is tightly coupled to pineal calcification. Using a dual threshold of |log₂ fold-change| > 1 under a Brownian-motion (BM) model and an FDR-adjusted P < 0.05 after Pagel’s λ transformation, it was found that every outlier is sharply up-regulated in the calcifying lineages, human, rat and goat, yet shows virtually no phylogenetic signal (λ = 1 × 10⁻⁷).

**Figure 4.**
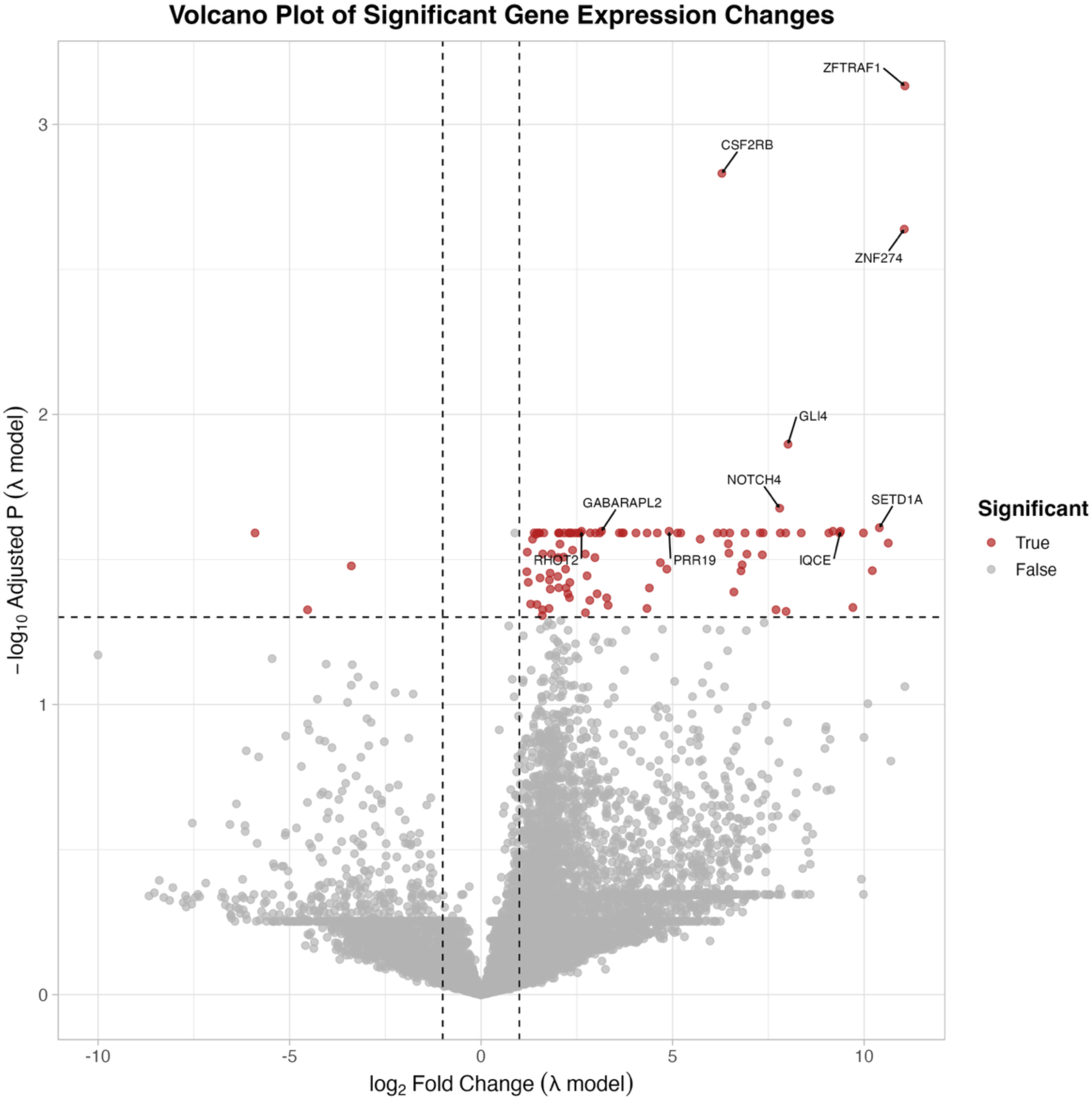
Volcano Plot of Significant Gene Expression Changes under the λ-Model Each point represents one gene tested by phylogenetic generalized least squares (PGLS) with a Brownian-motion transformation parameter λ estimated by maximum likelihood. The x-axis shows the log₂ fold change in expression (log₂FC) under the fitted λ-model; positive values indicate up-regulation, negative values down-regulation. The y-axis is –log₁₀ of the FDR-adjusted P-value (adj.Pλ), so that more statistically significant genes appear higher on the plot. Genes were pre-filtered to exclude any with missing logFCλ or Pλ estimates and those with model λ ≥ 0.7 (to avoid extreme phylogenetic signal). Significance was defined as |log₂FC| > 1 (i.e. at least a two-fold change) and adj.Pλ < 0.05. These criteria correspond to the vertical dashed lines at log₂FC = ±1 and the horizontal dashed line at –log₁₀(0.05) ≈ 1.3. Significant genes (meeting both thresholds) are colored red, whereas non-significant genes are colored grey.

**Figure 5.**
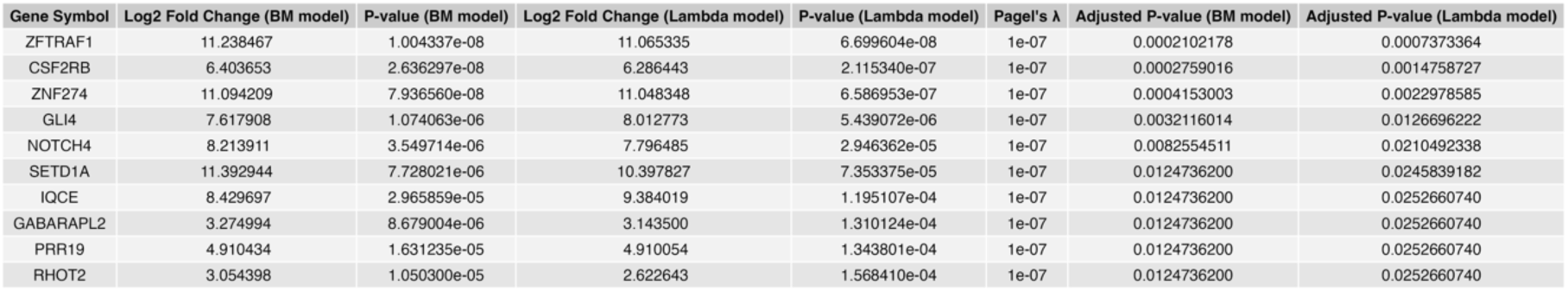
Top Ten Genes Significant in Both Brownian-Motion and λ-Transformed PGLS Models. This table lists the ten genes with the smallest FDR-adjusted P-values under the λ-model (adj.Pλ < 0.05), each of which also meets significance in the Brownian-motion (BM) model (adj.PBM < 0.05), exhibits absolute log₂ fold change > 1 in both models, and was fit with Pagel’s λ < 0.7. Columns include the official gene symbol; the log₂ fold change and nominal P- value from the BM model; the log₂ fold change and nominal P-value from the λ model; the maximum-likelihood estimate of Pagel’s λ (phylogenetic signal scaling factor); and the FDR- corrected P-values for each model. Genes are sorted by ascending adjusted P-value from the λ model. The table was rendered as a grid graphical object (gridExtra::tableGrob, rows suppressed) and exported at 18 × 4 inches, 300 dpi.

The most extreme responder is ZFTRAF1/CYHR1, whose effect size approaches twelve-fold under both BM and λ tests, mirroring its strong transcriptional induction in mineralizing human tissues.^44^ Two chromatin modifiers, SETD1A, a core H3K4 methyltransferase, and the SETDB1 recruiter ZNF274, follow closely, highlighting an epigenetic axis in the calcification program.^36,37^ Developmental signaling components also rise conspicuously: GLI4, a downstream effector of Hedgehog signaling, and NOTCH4, part of the Notch pathway implicated in vascular calcification, each exceed seven-fold change and remain significant after λ correction.^31,33^

Immune and stress cues are represented by CSF2RB, the common β-chain of GM-CSF/IL-3/IL-5 receptors, while cellular quality control is flagged by the autophagy mediator GABARAPL2 and the mitochondrial trafficking GTPase RHOT2/MIRO2.^34,39,40^ Rounding out the list are PRR19, recently linked to genome-maintenance during meiosis, and the cilia-anchoring Hedgehog regulator IQCE.^32,45,46^

Concordance between the BM and λ metrics, nearly identical log₂ fold-changes and sub-millimagnitude adjusted P-values, confirms that differential expression in these genes reflects calcification status rather than shared ancestry. Collectively they delineate a reproducible molecular signature of pineal mineralization that spans chromatin remodeling, developmental morphogenesis, cytokine signaling and organelle homeostasis.

## Discussion

The present comparative transcriptomic survey establishes the first phylogenetically controlled molecular portrait of pineal gland mineralization. By exploiting the natural contrast between three calcifying mammals (human, rat, goat) and three non-calcifiers that nevertheless share canonical melatonin circuitry (mouse, chicken, zebrafish), demonstrating that the appearance of corpora arenacea is accompanied by the concerted activation of a discrete, evolutionarily labile gene network rather than by stochastic, age-related precipitation. Species with brain sand segregated cleanly in principal-component space and shared a 103-gene “calcifier module” whose expression increase was independent of shared ancestry (Pagel’s λ ≈ 0). Conversely, the non-calcifier module was limited to three genes, RMI2, RASL11B, and GPR18, whose suppression in calcifiers hints at protective or housekeeping roles that are down-tuned once mineralization is underway.

Central to the calcifier module are developmental morphogen pathways, such as Hedgehog and Notch, that ordinarily orchestrate skeletal or vascular ossification.^33,47^ The marked up-regulation of GLI4, one of the GLI-family Hedgehog effectors whose paralogue GLI2 drives Runx2-dependent osteoblastogenesis, and of IQCE, which tethers the EVC-EVC2 complex to primary cilia to potentiate Hedgehog responses, points to a re-deployment of this embryonic axis in the adult pineal.^11,46^ Concurrently, the rise of NOTCH4, a Notch receptor that can trigger endothelial-to-smooth-muscle osteogenic crosstalk and calcification, suggests that Notch–Msx2 circuitry likewise contributes to acervulus formation.^48^ Comparative phylogenetic analysis of transcript abundance (Pagel’s λ < 0.1 calculated with the phylosignal framework) shows negligible phylogenetic signal for GLI4, IQCE and NOTCH4, indicating that their induction is tied to the calcified phenotype rather than shared mammalian ancestry.^49^

A second dominant theme is epigenetic re-patterning. Key chromatin regulators, including the H3K4 methyl-transferase SETD1A, the SETDB1-recruiting zinc-finger protein ZNF274, and the HUSH-complex stabilizer ATF7IP, sit at the center of this machinery.^37,38,50,51^ Taken together, these factors point to large-scale chromatin re-organization as a likely prerequisite for the transcriptional switch from neuro-endocrine to mineralizing states.^38,50^ Such a model meshes well with ultrastructural observations of concentric, bone-like lamellae in human acervuli and with the fact that successful osteogenic programs often require coordinated remodeling of activating H3K4 and repressive H3K9 marks.^1,51,52^ Future chromatin-profiling (e.g. CUT&RUN) experiments in rat and goat pinealocytes should therefore clarify whether SETD1A-dependent H3K4 trimethylation is indeed a gateway to mineral deposition, mirroring its documented importance in osteogenic differentiation.^51^

Inflammatory and cellular-stress cues also emerge as likely triggers of ectopic calcification. Pro-inflammatory cytokines (e.g., TNF-α, IL-6) are documented drivers of vascular and valvular calcification, implicating their downstream receptors in the process.^53^ CSF2RB encodes the common β-chain of the GM-CSF, IL-3 and IL-5 receptors, situating this gene at the heart of such cytokine signaling pathways.^54^ Autophagy- and mitochondria-quality-control genes reinforce this axis: GABARAPL2 is required for autophagosome–lysosome fusion, while RHOT2/MIRO2 coordinates mitochondrial trafficking to maintain organelle homeostasis. It is therefore suggested that sustained inflammatory signaling together with mitochondrial stress yields poorly cleared sub-cellular fragments that serve as nucleation sites for hydroxyapatite deposition. This view is underpinned by electron-microscopy evidence of calcium foci within degenerating pinealocyte vesicles in aged rats and by experiments showing that hypoxia, a canonical cellular-stress signal, accelerates vascular calcification.^7,55^

Two relatively understudied transcripts, PRR19 and ZFTRAF1 (also called CYHR1), warrant special attention in the context of pineal calcification. PRR19 partners with the cyclin-like factor CNTD1 and is required for timely repair of meiotic double-strand breaks (DSBs), forming crossover-specific recombination complexes.^45^ Because persistent DSB signaling can itself drive pathological mineralization in other tissues^56^, up-regulation of PRR19 may mark an active DNA-damage response during pineal stone formation. CYHR1, meanwhile, acts as a pro-survival factor in esophageal squamous-cell carcinoma, its knock-down suppresses proliferation, invasion, and tumor growth both in vitro and in vivo^44^; CYHR1’s function in endocrine contexts remains unexplored, leaving its pineal role unknown. Rat pineal organotypic cultures preserve viable, secretory pinealocytes that can be experimentally manipulated^57^, providing a tractable system in which RNA-interference or CRISPR approaches could determine whether PRR19 and CYHR1 are merely by-standers or active promoters of calcification.

### Limitations and future directions

A key limitation of the study is the reliance on a single high-quality transcriptome from each species, which prevents assessment of within-species variability as well as sex- or age-specific effects.^58^ Increasing biological-replicate depth, particularly in wild-type rodents, where rare micro-acervuli have been documented, should sharpen gene–phenotype correlations.^59^ Furthermore, applying single-nucleus RNA-seq and spatial transcriptomics will be essential to map the calcifier signature to defined pinealocyte subtypes or infiltrating immune cells and to reconstruct the temporal cascade that culminates in mineral nucleation.^60,61^ Finally, targeted perturbation of Hedgehog and Notch signaling, and of the SET1/SETD1A histone-methyltransferase that modulates clock-gene expression, could test whether disabling this program restores melatonin output and circadian-rhythm integrity.^62–64^

Together, these findings support the interpretation that pineal-gland calcification is a regulated, lineage-restricted trait driven by ectopic re-activation of developmental, inflammatory and epigenetic pathways rather than by passive precipitation^10,59,62^ The ten-gene core signature defined, analogous to transcriptional signatures that have guided mechanistic and therapeutic studies in other disease contexts^65^, provides a molecular foothold for dissecting the calcification program and for exploring interventions aimed at preserving pineal function, and thus circadian health, into old age.

